# Broad sensitivity of *Candida auris* strains to quinolones and mechanisms of resistance

**DOI:** 10.1101/2023.02.16.528905

**Authors:** Matthew B. Lohse, Matthew T. Laurie, Sophia Levan, Naomi Ziv, Craig L. Ennis, Clarissa J. Nobile, Joseph DeRisi, Alexander D. Johnson

## Abstract

The fungal pathogen *Candida auris* represents a severe threat to hospitalized patients. Its resistance to multiple classes of antifungal drugs and ability to spread and resist decontamination in health-care settings make it especially dangerous. We screened 1,990 clinically approved and late-stage investigational compounds for the potential to be repurposed as antifungal drugs targeting *C. auris* and narrowed our focus to five FDA-approved compounds with inhibitory concentrations under 10 µM for *C. auris* and significantly lower toxicity to three human cell lines. These compounds, some of which had been previously identified in independent screens, include three dihalogenated 8-hydroxyquinolines: broxyquinoline, chloroxine, and clioquinol. A subsequent structure-activity study of 32 quinoline derivatives found that 8-hydroxyquinolines, especially those dihalogenated at the C5 and C7 positions, were the most effective inhibitors of *C. auris*. To pursue these compounds further, we exposed *C. auris* to clioquinol in an extended experimental evolution study and found that *C. auris* developed only 2- to 5-fold resistance to the compound. DNA sequencing of resistant strains and subsequent verification by directed mutation in naive strains revealed that resistance was due to mutations in the transcriptional regulator *CAP1* (causing upregulation of the drug transporter *MDR1*) and in the drug transporter *CDR1*. These mutations had only modest effects on resistance to traditional antifungal agents, and the *CDR1* mutation rendered *C. auris* more sensitive to posaconazole. This observation raises the possibility that a combination treatment involving an 8-hydroxyquinoline and posaconazole might prevent *C. auris* from developing resistance to this established antifungal agent.

**Abstract Importance:** The rapidly emerging fungal pathogen *Candida auris* represents a growing threat to hospitalized patients, in part due to frequent resistance to multiple classes of antifungal drugs. We identify a class of compounds, the dihalogenated hydroxyquinolines, with broad fungistatic ability against a diverse collection of 13 strains of *C. auris*. Although this compound has been identified in previous screens, we extended the analysis by showing that *C. auris* developed only modest 2- to 5-fold increases in resistance to this class of compounds despite long-term exposure; a noticeable difference from the 30- to 500- fold increases in resistance reported for similar studies with commonly used antifungal drugs. We also identify the mutations underlying the resistance. These results suggest that the dihalogenated hydroxyquinolines are working inside the fungal cell and should be developed further to combat *C. auris* and other fungal pathogens.

**Tweet:** Lohse and colleagues characterize a class of compounds that inhibit the fungal pathogen *C. auris*. Unlike many other antifungal drugs, *C. auris* does not readily develop resistance to this class of compounds.

## Introduction

*Candida auris* is a rapidly emerging multidrug resistant pathogen responsible for invasive fungal infections in hospitalized patients. Similar to other *Candida* species, *C. auris* predominately induces candidemia in the immunocompromised and those subjected to prolonged hospitalization in ICU wards or long-term care facilities. In untreated patients, invasive candidemia has a mortality rate of approximately 60%, which only improves to approximately 40% with antifungal therapy (1, 2). The threat of *C. auris* is compounded by persistent colonization in previously exposed patients and pervasive spread through hospital wards despite multiple rounds of decontamination (3, 4). Furthermore, *C. auris* can spread through long-term care and skilled nursing facilities with older and ventilator-dependent patients being especially at risk for infection; several *C. auris* outbreaks associated with COVID-19 treatment units have been reported (5–9). This pervasive colonization, in combination with frequent resistance to one, two, or even all three major classes of antifungals, makes *C. auris* a growing threat to our most vulnerable patients (10). For these reasons, the World Health Organization’s recently released fungal priority pathogens list includes *C. auris* as one of four fungal pathogens in the critical (as opposed to high or medium) priority group (11).

*C. auris* represents a relatively new threat to hospitalized patients. It was first reported in Japan in 2009 (12) and was subsequently found to have five clades (I-V) that localize to distinct geographic locations (13, 14). The five clades, which are geographically represented by South Asia (I), East Asia (II), Africa (III), South America (IV), and Iran (V), have different frequencies of antifungal resistance and two distinct mating types (13–16). Clades I, III, IV, and V have been linked to outbreaks of invasive infections while clade II is more commonly associated with ear infections (17, 18). While specific clades typically predominate in different parts of the world, the US, Canada, UK, and Kenya have identified infections associated with a range of isolates from multiple clades (16, 19–21). Nearly all *C. auris* isolates are highly resistant to fluconazole; more than half are resistant to voriconazole; a third are resistant to amphotericin B; and some isolates are resistant to all three major classes of antifungal drugs including the echinocandins caspofungin and micafungin (16, 22–26). Given the high mortality, limited treatment options, and growing threat to vulnerable patient populations, there is an urgent need to develop new antifungal agents to combat *C. auris*.

Several approaches have been taken to identify new antifungal agents effective against *C. auris* (for more detail on this topic and the current state of the antifungal drug development pipeline, see (27–33)). The most straightforward approach has focused on the evaluation of the effectiveness of the newest members of common antifungal classes (e.g. the echinocandin rezafungin (CD101)) (34, 35). A closely related line of investigation has focused on testing the lead compound(s) from new classes of potential antifungal agents in development and/or in the clinical testing pipeline (e.g. the fungal inositol acylase inhibitor fosmanogepix/APX001 (36–38) and the glucan synthesis inhibitor ibrexafungerp/SCY-078 as well as the second generation fungerp analog SCY-247 (39–43)). A broader approach, and one less dependent on existing antifungal drug development pipelines, involves screening libraries of FDA-approved compounds and/or drug like compounds to repurpose existing clinical compounds for use against *C. auris*. These types of screens have identified a number of promising compounds, including ebselen, miltefosine, and alexidine dihydrochloride. Three of these published screens were conducted with the same library (the Prestwick Chemical Library with 1,280 compounds), but the hits from these screens were not always concordant (49 different compounds were identified in these screens: 3 in all three, 18 in two, and 28 in only one) (44–49).

To identify potential drug repurposing candidates, we conducted a broad primary drug screen that examined the effect of 1,990 clinically approved or late-stage investigational compounds on three *C. auris* strains from the clades most associated with invasive infections (I, III, and IV). From the 86 candidate compounds identified during this preliminary screen, we found five FDA-approved compounds with half-maximal inhibitory concentrations (IC50) that were less than 10 µM for *C. auris* and at least an order of magnitude lower in toxicity to three human cell lines. For one of these compounds, the 8-hydroxyquinoline clioquinol, we identified mutations conferring resistance by growing two *C. auris* isolates for an extended period in the presence of the compound, followed by whole genome sequencing. To prove that the mutations we identified caused the resistance, we reconstructed the mutations in a naïve strain (using CRISPR-based approaches) and showed they conferred resistance.

## Results

### The hydroxyquinolines broxyquinoline, chloroxine, and clioquinol inhibit C. auris growth at submicromolar concentrations

A primary screen of 1,990 compounds, consisting of clinically approved drugs, late-stage investigational compounds, and drug-like compounds, was performed using three strains of *C. auris* that were selected based on their range of resistance to the three main classes of antifungal agents and to represent the three clades most associated with invasive infections (Figure 1A). Strain AR-387 (B8441/MLY1543) originated in Pakistan, belongs to the South Asian clade, and is sensitive to fluconazole, caspofungin, and amphotericin B. Strain AR-386 (B11245/MLY1542) originated in Venezuela, belongs to the South American clade, is sensitive to both caspofungin and amphotericin B but resistant to fluconazole. Strain AR-384 (B11222/MLY1540) originated in South Africa, belongs to the African clade, is sensitive to amphotericin B and resistant to both fluconazole and caspofungin.

**Figure 1.**
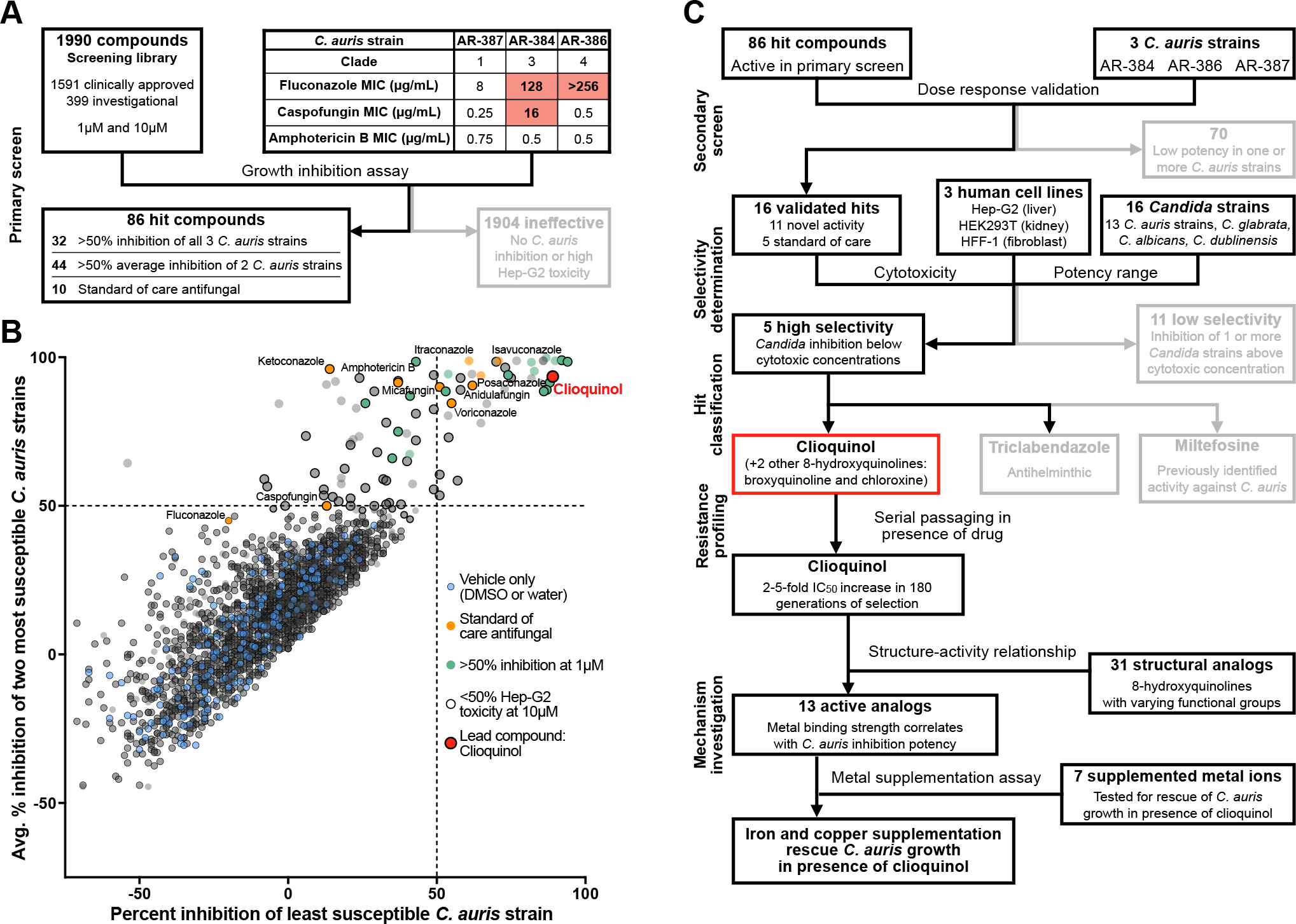
A screen of 1,990 clinically approved and investigational compounds for *in vitro* inhibition of *C. auris* identified 86 candidates for further evaluation. (A) Workflow for the primary screening of 1,990 clinically approved and investigational compounds for *in vitro* inhibition of three *C. auris* strains; compounds were screened at both 1 µM and 10 µM. (B) The percent inhibition relative to untreated controls (DMSO or water alone) for each compound at 10 µM. For each compound, the lowest percent inhibition achieved against any of the three screened *C. auris* strains is plotted on the x-axis and the average percent inhibition against the other two strains is plotted on the y-axis. Compounds in the upper left quadrant effectively inhibited two of the three *C. auris* strains; compounds in the upper right quadrant effectively inhibited all three strains. (C) The secondary screening pipeline for the 86 compounds identified in the initial screen for their ability to inhibit at least two of the three *C. auris* strains by at least 50 percent. Additional screening monitoring inhibition of additional *C. auris* strains, other *Candida* species, and lack of toxicity to human cells yielded 5 highly selective candidates (see Figure 2). Figures 3 and 4 detail additional criteria depicted in the flow chart.

These three strains were screened for growth inhibition at two concentrations (1 µM and 10 µM) of drugs from the Selleck Chem FDA-Approved Drug Library (#L1300, 1,591 compounds) and the Medicines for Malaria Venture Pandemic Response Box (399 compounds) (Figure 1A). Compounds with a B-score (a non-control based method accounting for systematic errors including plate position effects (50, 51)) greater than 0.1 and greater than 50 percent inhibition (measured by OD600) were considered initial hits (Figure 1B, File S1). The 86 initial hits include 10 established standard-of-care compounds for fungal infections, 44 compounds active against only two of the three *C. auris* strains tested, and 32 compounds active against all three (Figure 1B, File S1). A secondary screen was performed to confirm the antifungal activity observed in the primary screen and to estimate the half maximal inhibitory concentrations (IC50s) for these compounds (Figure 1C). All 86 compounds were screened at 8 concentrations ranging from 0.3 to 100 µM against the same three strains (AR-387, AR-386, and AR-384) (Figure 2A, File S2). We then selected 11 compounds that had estimated IC50s less than 10 µM in this secondary screen and did not belong to the three main known classes of antifungal drugs. Three additional compounds (sirolimus, everolimus, temsirolimus) met these criteria, but were not selected for further investigation because of their known immunosuppressive activity.

**Figure 2.**
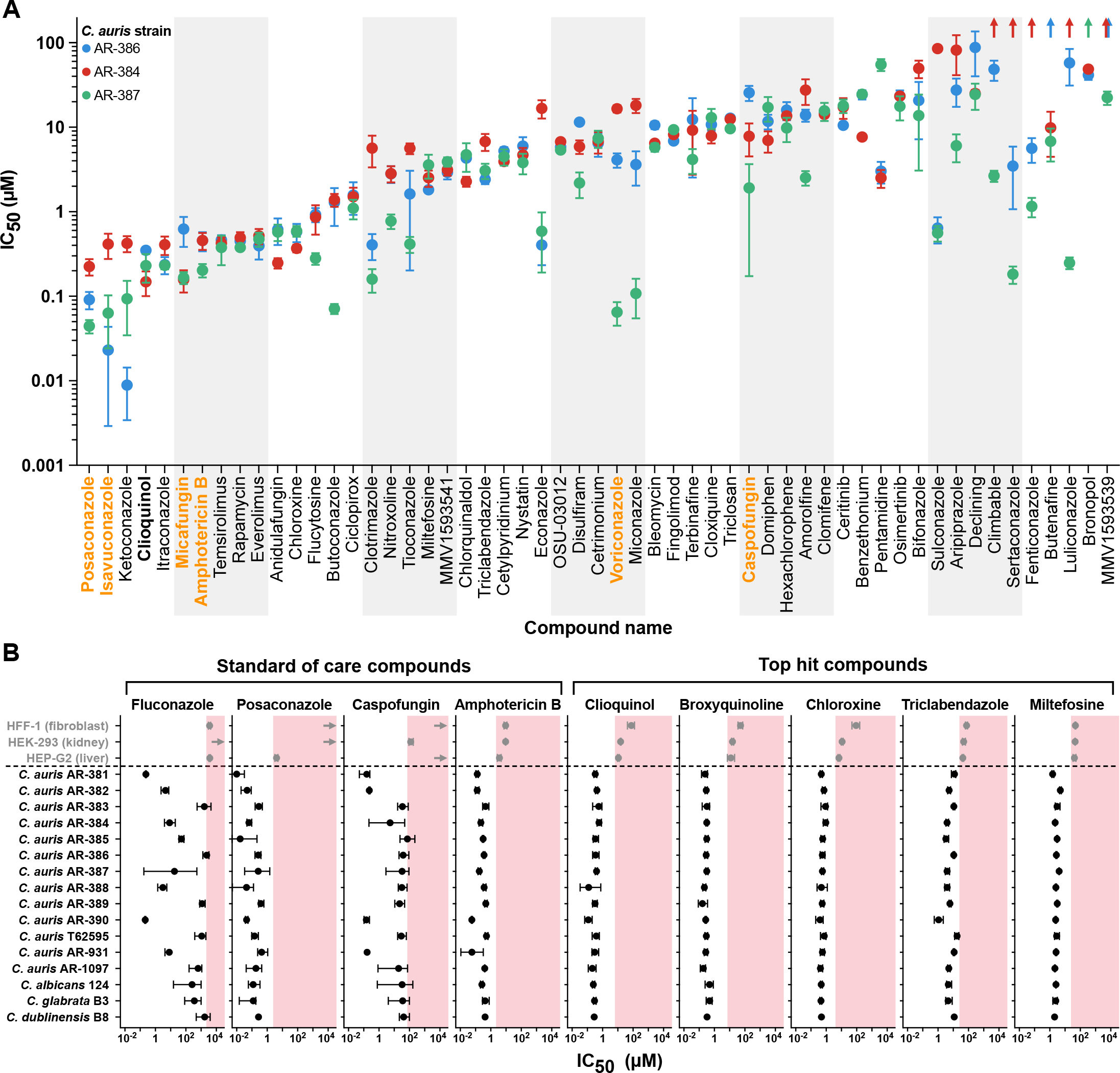
Secondary screening and selectivity measurement identified five promising compounds, including three hydroxyquinolines. (A) In the secondary screen, the results with 31 of the 86 hits identified in the primary screen did not repeat and were eliminated from further consideration. IC50 values against three *C. auris* strains for the remaining 55 compounds were calculated from 8-point dose-response curves. Points represent the mean of three biological replicates, error bars represent the standard error of the mean, and upwards arrows indicate IC50 values greater than the highest concentration tested, 100 µM. Representative standard-of-care compounds used to treat fungal infections are indicated in orange. (B) IC50 values for 4 standard-of-care drugs and 5 finalists from the screen against 13 *C. auris* strains, strains from 3 additional *Candida* species, and 3 human cell lines. Pink shaded regions mark concentrations above the lowest observed IC50 for that compound against a human cell line. Values represent the mean of three biological replicates, error bars represent the standard error of the mean, and right-pointing arrows indicate IC50 values greater than 1000 µM.

An optimal drug candidate would have antifungal activity against a wide range of *C. auris* isolates at concentrations that are not toxic to human cells. To determine if any of the 11 selected compounds fulfilled these criteria, IC50s were determined for 16 *Candida* strains, including 13 *C. auris* strains, covering all five clades, and one strain each of *Candida albicans*, *Candida glabrata*, and *Candida dubliniensis*. The IC50s of these 11 compounds were also determined for three common human cell lines, HEK293 (kidney), HEPG2 (liver) and HFF1 (fibroblast) (Figure 1C). Of the 11 compounds, five had IC50s that were at least 10-fold less than their lowest IC50 for human cells, suggesting the possibility of a therapeutic window (Figure 2B). These five compounds included three hydroxyquinolines, broxyquinoline, chloroxine, and clioquinol, and two anti-protozoals, miltefosine and triclabendazole (Figure 2B). Unlike many of the traditional antifungal agents, there was minimal variation in the IC50s observed for miltefosine, broxyquinoline, chloroxine, and clioquinol across all 13 *C. auris* strains tested (Figure 2B). These results are broadly consistent with previous repurposing screens, which have reported effectiveness by miltefosine (46), several different hydroxyquinolines including chloroxine and clioquinol (45–47, 49, 52–54), as well as pentamidine (one of the six compounds that performed poorly in our human cell toxicity tests) (44, 46).

### C. auris develops only moderate resistance to clioquinol despite extended exposure

*C. auris* has repeatedly demonstrated a propensity for rapid acquisition of resistance when exposed to antifungal drugs. Indeed, experimental evolution studies have produced 30- to more than 500-fold increases in resistance to fluconazole or caspofungin in as few as two or three 24- or 48-hour passages (55–57). To evaluate the ability of *C. auris* to develop resistance to clioquinol, and to assess the resulting determinants of resistance, two independent cultures of *auris* AR-384 (discussed above) and one culture of AR-390, another South Asian clade I strain with greater resistance to fluconazole and amphotericin B than AR-387, were serially passaged 30 times (roughly 150-200 generations) in the presence of increasing concentrations of clioquinol, ranging from 0.4-0.75 µM at the start to 2-8-4.4 µM at the end (Figure 1C). By the end of the drug selection, the clioquinol IC50 increased 2.4- and 5.2-fold relative to parental strains (from 0.75 µM to 1.80µM and from 0.29 µM to 1.48µM) in the AR-384 and AR-390 backgrounds, respectively (Figures 3A, 3B). The IC50 increases occurred over multiple steps, showed some variation in the rate of resistance evolution, and leveled off around passage 16 for each culture (Figure 3A). Notably, the degree of resistance arising from more than two months of exposure to clioquinol was less than has been observed for fluconazole or caspofungin over shorter time frames. Furthermore, the clioquinol-evolved strains remained sensitive to clioquinol concentrations at least 5-fold below the minimal toxic concentrations observed for human cells.

**Figure 3.**
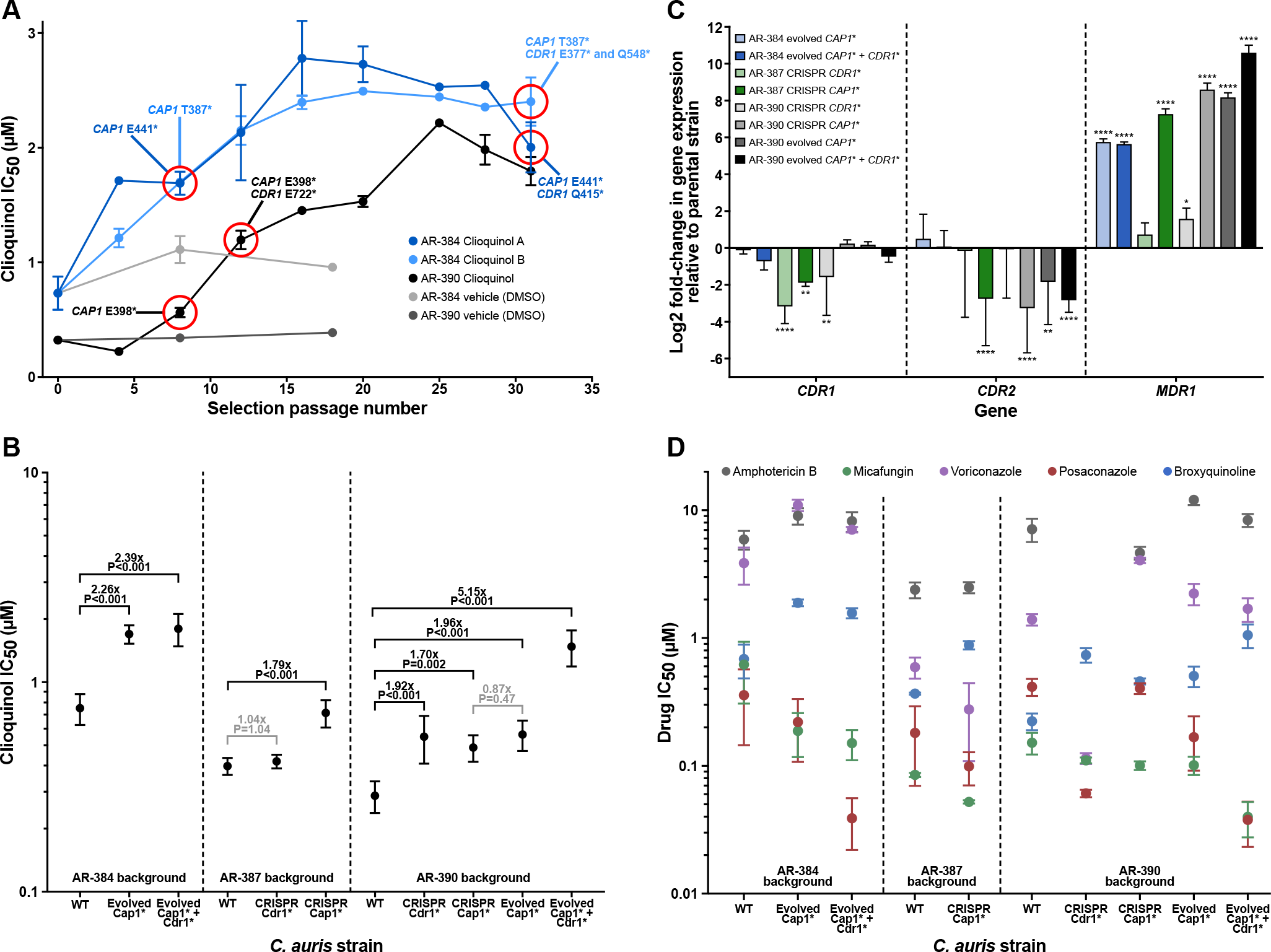
*C. auris* developed 2- to 5- fold resistance to clioquinol in an extended evolution experiment due to mutations in the transcriptional regulator *CAP1* and the *CDR1* drug pump. (A) Clioquinol IC50 against *C. auris* isolates that were selected in the presence of clioquinol. Parallel cultures were grown with serial passaging of strains AR-384 (blue, two independent cultures) and AR-390 (black, one culture) in the presence of clioquinol or vehicle control (DMSO; grey). Whole genome sequencing was performed for a subset of *C. auris* passage populations or single cells isolated from the cultures and selected mutations identified are indicated. (B) Comparison of clioquinol’s IC50 against parent, evolved, and CRISPR/Cas9 engineered *C. auris* mutant strains from three different backgrounds. The fold-changes in IC50 between relevant related strains are shown; the statistical significance of observed differences as determined with unpaired t-tests (two-tailed, equal variance) is indicated. (C) Changes in expression of the *CDR1*, *CDR2*, and *MDR1* drug pump genes in evolved and CRISPR/Cas9 engineered *C. auris CAP1* and *CDR1* mutant strains from three different backgrounds in the absence of clioquinol. Gene expression changes are shown as the Log2 fold-change relative to the parental (WT) strain for that background. Statistical significance was determined with unpaired t-tests (two-tailed, equal variance, * P<.05; ** P<.01; *** P<.001; **** P<.0001). (D) Comparison of IC50 values for 5 other drugs against parent, evolved, and CRISPR-Cas9 engineered mutant *C. auris* strains. Values represent the mean of three independent experiments; error bars represent the standard error of the mean.

#### Mutations in the genes CAP1 and CDR1 arose during extended clioquinol exposure and are the cause of resistance

The resistance of the evolved strains persisted when these strains were regrown for several days in the absence of clioquinol, indicating that resistance was linked to one or more genome mutations rather than a reversible, physiological response to clioquinol. To test this hypothesis and identify the determinant(s) of clioquinol resistance, we performed whole genome sequencing on two single colony isolates from the endpoint of the resistance experiment for each of the three cultures as well as from populations harvested from selected intermediate points. We observed premature termination mutations in the C-terminal end of the transcriptional regulator *CAP1* (B9J08_005344 / CJI97_005427) by the eighth passage in all three cultures (16 days growth or approximately 35 to 40 generations) (Figure 3A). Each mutation was distinct (E398* in AR-390; either E441* or a 8-bp deletion resulting in T387* in AR-384) and were in 76 to 98% of the population sample reads at passage eight. The same *CAP1* mutations were in 96 to 98% of population sample reads and in all six single cell samples at the endpoint of the experiment. *CAP1* mutations were not observed in the DMSO treated control cultures that were grown and sequenced in parallel. Based on additional experiments described below, we believe these truncation mutations produce hyperactive Cap1 proteins.

Sometime after the eighth passage (by passage 12 in AR-390 and between passages 8 and 31 in AR-384), additional mutations arose in the ATP-binding cassette (ABC) drug transporter *CDR1* (B9J08_000164 / CJI97_000167). The mutations in this gene were different in the three cultures, and likely result in loss of function of the gene (see below; E722* in AR-390; K909N or Q415* in one AR-384 culture, and Q548* or an 8bp deletion whose resulting frame shift affected 34 different amino acids at aa343-376 before a premature stop codon at aa377 in the other AR-384 culture) (Figure 3A). These *CDR1* mutations were not as prevalent in the populations as the *CAP1* mutations, comprising between 25 and 97% of reads in population samples and were observed in only four of the six single-cell endpoint samples. No *CDR1* mutations were observed in the DMSO-treated control cultures. We note that the apparent selective pressure to inactivate the *CDR1* drug pump suggests that hydroxyquinolines function against *C. auris*, at least in part, at the cell surface or inside the cell rather than by chelating soluble iron, copper, or zinc in the media, consistent with a previous report that hydroxyquinolines sequester metals in *Saccharomyces cerevisiae’s* plasma membrane (58). It may seem paradoxical that inactivating a drug pump results in increased resistance − rather than sensitivity − to clioquinol; however, in addition to exporting drugs, *CDR1* has been implicated in the translocation of phosphoglycerides from the internal to the external plasma membrane (59, 60). In principle, the hydroxyquinolines could be affecting another function of Cdr1 instead of, or in addition to, exporting drugs from the cell. It is also possible that Cdr1 functions to import hydroxyquinolines into the cell.

To determine whether the *CAP1* or *CDR1* mutations were indeed causal for increased clioquinol resistance, the *CAP1* E398* and the *CDR1* E722* mutations were introduced by CRISPR-Cas9 gene editing into the parental strains AR-390 and AR-387. The *CAP1* E398* mutation caused an increase in resistance of 1.7- to 1.8-fold to clioquinol in both strains (Figure 3B), similar to that observed for the evolved *CAP1*\* strains, indicating that the *CAP1* mutation is the cause of resistance in the evolved strains. Introducing the *CDR1* E722* mutation did not significantly affect clioquinol resistance in AR-387 (nor did deletion of *CDR1*) but it did increase resistance 1.9-fold in AR-390 (Figure 3B). Thus, the *CAP1* truncation (which results in a gain-of-function mutation, see below) can explain much of the clioquinol resistance that arose during the experimental evolution; the subsequent *CDR1* mutation (which likely results in a loss-of-function) could account for the smaller resistance increases observed later in the experiment.

### *CAP1* truncation results in increased *MDR1* expression

Cap1 is a transcriptional regulator. In *C. albicans*, it promotes expression of the major facilitator superfamily (MFS) drug transporter *MDR1,* and it has been reported that C-terminal truncations of *CAP1*, like those we isolated from our drug-resistance screen, cause a hyperactive phenotype which increases *MDR1* expression and thereby increases fluconazole resistance (61–64). To test whether this is also the case with our resistant *C. auris CAP1* mutants, we used RT-qPCR to examine the levels of *MDR1* expression along with two other drug transporters: *CDR1* and *CDR2*. We found that *MDR1* expression had increased more than 50-fold by passage 8 in the AR-384 background compared to the starting strain and remained at this level at the end of the experiment (Figure 3C). In the AR-390 background, where baseline expression began roughly 30-fold lower than AR-384, *MDR1* expression increased nearly 300- fold compared to the starting strain by passage 8 and over 1500-fold by the final selection passage, reaching a similar level to the evolved AR-384 strains (Figure 3C). In contrast to *MDR1*, expression of *CDR1* and *CDR2* changed only minimally during the course of the selection experiment (1.7-fold down and 1.1-fold up in AR-384, 1.4-fold down and 7-fold down in AR-390, respectively) (Figure 3C). We note that we profiled transcript levels in the cells in the absence of clioquinol; thus, the changes we observed are due to the mutations and are not dependent on the presence of the compound.

To verify these results, transcript levels were also assessed in the mutant stains constructed in the AR-387 and AR-390 strain backgrounds, again in the absence of clioquinol. Consistent with the results for the evolved strains, the replacement of the wildtype *CAP1* with the *CAP1* E398* mutation resulted in minimal expression change for *CDR1* and *CDR2*, but 150- and 390-fold increases in *MDR1* expression in the AR-387 and AR-390 strain backgrounds, respectively (Figure 3C). The results of these experiments show that *MDR1* expression is significantly increased by truncation mutations in *CAP1* both in the evolved strains and the genetically modified strains.

#### Extended clioquinol exposure has only modest effects on resistance to traditional antifungal agents

The *CAP1* truncation mutations that arose in response to clioquinol could plausibly produce resistance to the existing antifungal agents used to treat *C. auris* infections.

Conversely, given the role of *CDR1* in azole resistance (48, 65–69), the *CDR1* mutations that arose in our drug-resistant cultures could increase sensitivity to existing antifungal agents. To test this hypothesis, the sensitivities of the evolved strains to a second hydroxyquinoline, broxyquinoline, and to the traditional antifungal agents voriconazole, posaconazole, micafungin, and amphotericin B were determined. As expected, the modest clioquinol resistance was also observed for the structurally similar broxyquinoline (Figure 3D). Clioquinol resistance had only a subtle, if any, effect on sensitivity to amphotericin B, modestly increased susceptibility to micafungin (four-fold), and modestly increased (less than three-fold) resistance to the azole voriconazole (Figure 3D). An unexpected property of the evolved strains containing the *CDR1* loss-of-function mutation was its 10-fold greater sensitivity to the azole posaconazole (Figure 3D). These findings indicate that, for *C. auris*, mutations resulting from long-term exposure to clioquinol will not necessarily result in increased resistance to commonly used antifungal agents. Indeed, these observations suggest that combining an 8-hydroxyquinoline with posaconazole might leave *C. auris* especially vulnerable to the latter.

#### A variety of dihalogenated hydroxyquinolines inhibit C. auris

As described above, three hydroxyquinolines with submicromolar antifungal activity against *C. auris* were identified in our screen. To evaluate the structure-activity relationship (SAR) of this class of compounds, the activities of 32 quinoline derivatives (29 compounds plus clioquinol, broxyquinoline, and chloroxine, the three identified in the screen) were tested against three strains of *C. auris* (AR-387, AR-384, AR-386) (Figure 1C). Seventeen of these compounds showed activity against all three *C. auris* strains tested. All active compounds except for one [1,10-phenanthroline monohydrate] were hydroxyquinolines, which have a hydroxide at C8 position (Figure 4A, File S3). Five compounds, including clioquinol, broxyquinoline, and chloroxine, had an IC50 less than 1µM; all five of these compounds were dihalogenated at the C5 and C7 positions, suggesting the importance of these modifications in antifungal activity.

**Figure 4.**
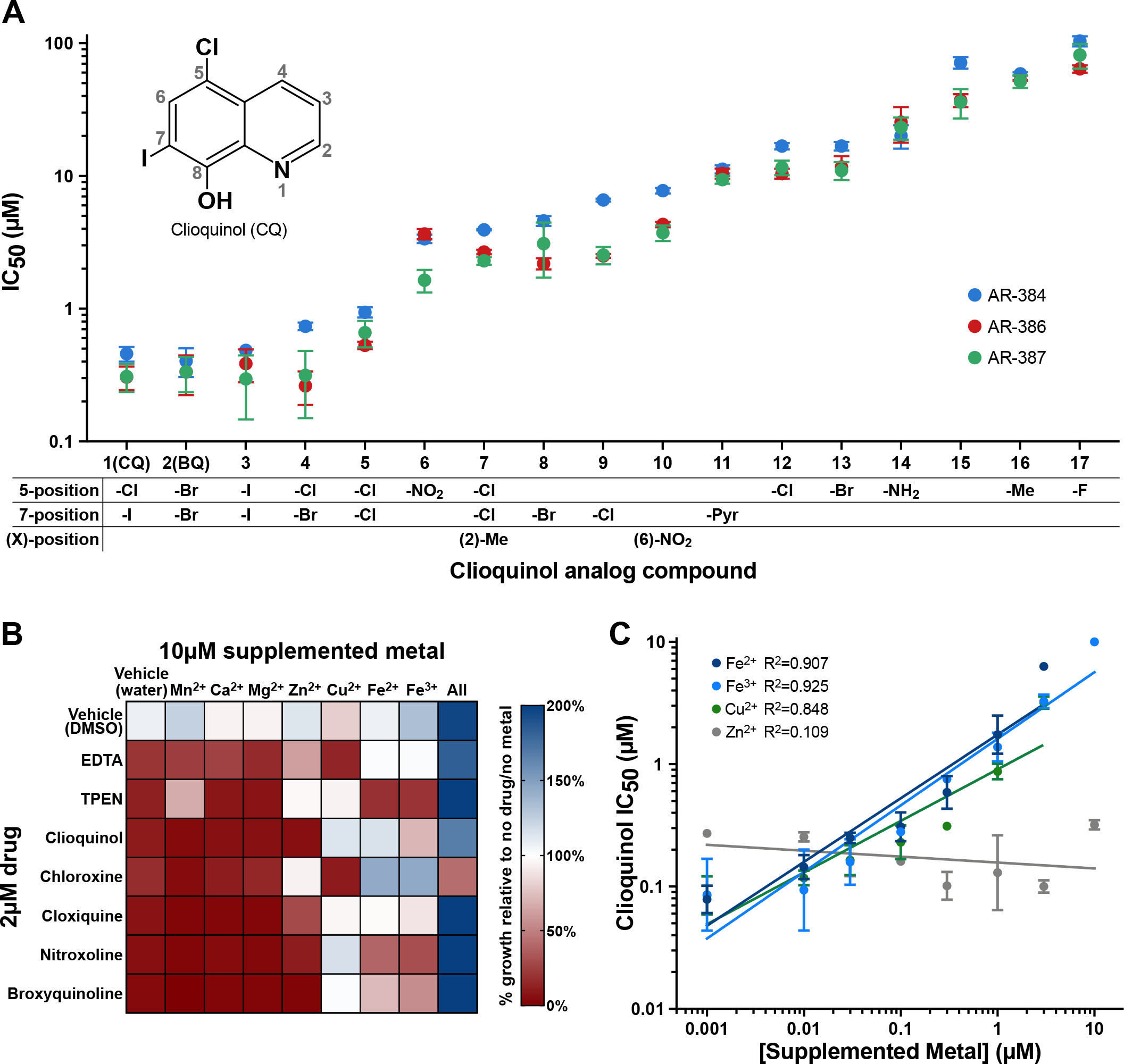
Out of the 32 SAR compounds (including clioquinol), dihalogenated 8- hydroxyquinolines were the most effective inhibitors of *C. auris*, and their activity is mitigated by excess iron or copper in the growth medium. (A) IC50 of clioquinol and 16 structural analogs against three *C. auris* strains. Clioquinol (upper left inset) is comprised of a quinoline core with a 5-position chlorine, 7-position iodine, and 8-position hydroxy group, the functional groups at these and other positions are indicated for each of the structural analogs. A further 15 clioquinol analogs (not shown) were tested and lacked activity against *C. auris*. (B) Heat map of *C. auris* growth (with and without 8-hydroxyquinoline compounds) in the presence of 10 µM metal ions. The effects of metal chelators EDTA and TPEN were also analyzed. The 8- hydroxyquinolines and the metal chelators inhibited growth (red on heat map) and addition of iron or copper mitigated these effects in at least some strains. (C) Relationship between supplemented metal concentration and clioquinol IC50. The linear regression line is shown for each metal. The points represent the mean of three independent experiments; error bars represent the standard error of the mean.

Compounds with single halogen or other chemical groups at either the 5- or 7-positions generally had an IC50 in the 2 to 20 µM range, only four of the 20 compounds with a hydroxide at C8 position lacked activity against *C. auris*. Eleven of the twelve compounds without an 8- position hydroxy group tested lacked activity against *C. auris*. (Figure 4A, File S3). We note that these trends are largely consistent with those previously reported for halogenated hydroxyquinoline activity against several fungal species, none of which were *Candida* (70–72).

Previous studies (including transcriptional profiling, enzyme activity assays, and cellular metal abundance quantification using inductively coupled plasma mass spectrometry) showed that *S. cerevisiae* cells treated with clioquinol behave as if they are starved for iron, copper, and zinc. These metals are sequestered in the plasma membrane and depleted in the cytosol of *S. cerevisiae* cells exposed to clioquinol (58, 73, 74). To investigate this effect with respect to *C. auris*, clioquinol treated cells were grown in the presence of excess iron, copper, or zinc (Figure 1C). The addition of excess iron (either Fe^2+^ or Fe^3+^) or, to a lesser extent, copper, to clioquinol- treated media mitigated the inhibitory effects of the drug and restored growth (Figures 4B, 4C). Increasing concentrations of Cu^2+^, Fe^2+^, or Fe^3+^ strongly correlated with increased clioquinol IC50, indicating that the increase in *C. auris* viability is concentration dependent (Figure 4C).

Increasing concentrations of Zn^2+^, on the other hand, had no effect on clioquinol IC50 (Figure 4C). Addition of excess iron or copper also restored growth in the presence of several other 8- hydroxyquinolines (Figure 4B). These effects were independent of the order of addition: either introducing iron to *C. auris* cells that had been pretreated with clioquinol for 21 hours or introducing metals to clioquinol-treated media prior to *C. auris* inoculation permitted growth (Figures 4B, S1A).

We note that the reversal of the inhibitory effects of hydroxyquinolines by exogenously added metals could be due, at least in part, to simply lowering the free concentration of hydroxyquinolines in the media. In this regard, the concentration of metals needed to overcome the inhibitory effects are in the same range as the concentrations of hydroxyquinolines needed to inhibit *C. auris*. The exact mechanisms by which hydroxyquinolines inhibit *C. auris* growth remain to be determined. It is a plausible hypothesis that the inhibitory effects arise from metal sequestration within the cell, perhaps in the fungal plasmid membrane as observed in *S. cerevisiae* (58). This idea is consistent with our observations that drug pump expression plays a role in the mechanism of resistance.

#### Clioquinol is fungistatic to C. auris

Previous reports have reached different conclusions as to whether clioquinol is fungistatic or fungicidal against *Saccharomycotina* species such as *S. cerevisiae* or *C. albicans* (75, 76). We quantified, by plating assays, the viability of *C. auris* cells treated with clioquinol for 22 hours and found that viability changed little relative to DMSO-treated controls (Figure S1B).

As such, we conclude that clioquinol is fungistatic, rather than fungicidal, to *C. auris*, at least in the 22-hour time frame examined.

## Discussion

*C. auris* represents a growing threat to hospitalized patients due to (1) the inherent resistance of many strains to one or more of the three major classes of antifungal drugs used in the clinic, (2) the propensity for sensitive *C. auris* strains to rapidly develop resistance, and (3) its ability to spread and resist decontamination in healthcare settings. We screened 1,990 compounds for the potential to be repurposed as antifungal agents targeting *C. auris*. Among the most promising hits from this screen were the dihalogenated hydroxyquinolines broxyquinoline, chloroxine, and clioquinol. A further structure-activity relationship study identified two additional, related compounds. Dihalogenated hydroxyquinolines have been identified in other small molecule screens for activity against *C. auris*, and we pursued this class of compounds further.

The five hydroxyquinolines mentioned above had IC50s less than 1 µM and, are especially interesting candidates for repurposing. First, we observed little variation in the IC50s of individual hydroxyquinolines across the variety of clinical isolates we tested (13 strains, including representatives of all five known clades), suggesting there is little inherent resistance to hydroxyquinolines. Second, although the dihalogenated hydroxyquinolines were the most efficient hydroxyquinolines in our SAR screen, a wide range of chemical space in this family remain to be explored (e.g. different side groups at C2 and different active groups replacing the halogen at C5 or C7). Third, several hydroxyquinolines have (or have had) both topical and oral forms (for use against dandruff and scalp dermatitis, eczema, fungal skin infections, and infectious diarrhea caused by protozoa or Shigella bacteria) suggesting that hydroxyquinolines might be effective against both internal and skin-based infections. Finally, the hydroxyquinolines are of interest because of the limited degree to which *C. auris* developed resistance, despite long-term exposure in our experiments. The roughly 2- to 5-fold increases in resistance we observed remain well below the minimum toxicities seen for human cells and are significantly less than the 30- to 500- fold increases in resistance reported for similar studies with fluconazole or caspofungin, two of the most widely used antifungal drugs (55–57). Furthermore, hydroxyquinoline resistance resulted in only modest, if any, increases in resistance to other antifungal agents, and it led to *increased* sensitivity to at least one antifungal agent in common use, posaconazole. The possibility exists that a combination treatment involving a hydroxyquinoline and posaconazole might leave *C. auris* trapped between increasing *CDR1* expression to resist posaconazole, becoming more sensitive to the hydroxyquinoline, or decreasing *CDR1* expression to resist the hydroxyquinoline, becoming more sensitive to posaconazole, a hypothesis that is amenable to testing in animal models and in culture.

When considering repurposing the hydroxyquinolines as a treatment for *C. auris*, it is important to note that an oral version of clioquinol was withdrawn from usage as an antiparasitic in the early 1970s following a report associating it with an outbreak of subacute myelo-optic neuropathy (SMON) in Japan (77–80). Since that time, however, the data in the report linking SMON and clioquinol has been questioned; it has been noted that similar associations did not occur in countries with higher clioquinol usage at the time and that many of the patients developing SMON had not taken clioquinol prior to the onset of symptoms (81, 82). As such, there has been recent interest in using clioquinol to treat both Alzheimer’s disease and cancer (58, 74, 77, 78, 83–86). The work reported here suggests that clioquinol and its derivatives could also be developed as effective antifungal agents, particularly against the emerging pathogen *C. auris*. Consistent with this, we note that the related hydroxyquinoline nitroxoline (which had an IC50 of 1.6-3.6 µM against *C. auris* in our SAR studies) has recently been reported to be effective at treating granulomatous amebic encephalitis caused by the amoeba *Balamuthia mandrillaris* (87).

## Methods

### Drug Libraries

The primary screen in this study used both the Selleck Chem FDA-Approved Drug Library (#L1300, 1,591 compounds) and the Medicines for Malaria Venture Pandemic Response Box (399 compounds). Details about the sources of drugs for subsequent assays can be found in File S4.

### Media

Cells were allowed to recover from glycerol stocks for at least two days at 30°C on yeast extract peptone dextrose (YEPD) plates (2% Bacto^TM^ peptone, 2% dextrose, 1% yeast extract, 2% agar). Unless otherwise noted, overnight cultures for assays, recovery dilutions, and assays were performed in RPMI-1640 media (containing L-glutamine and lacking sodium bicarbonate, MP Biomedicals #0910601) supplemented with 34.5g/L MOPS (Sigma, M3183) and adjusted to pH 7.0 with sodium hydroxide before sterilizing with a 0.22 µm filter.

### Strains

A full list of strains used in this study can be found in File S4. Twelve of the thirteen *C. auris* isolates used in this study were acquired from the Centers for Disease Control and Prevention’s Antibiotic Resistance Isolate Bank *Candida auris* panel (https://wwwn.cdc.gov/ARIsolateBank/Panel/PanelDetail?ID=2). The remaining *C. auris* isolate was a previously reported isolate from a patient at UCSF; this strain is a member of the South Asian clade that has some resistance to fluconazole (MIC 32 µg/mL) and was provided by the UCSF Clinical Laboratories at China Basin (88). For *C. albicans* (SC5314), *Candida dubliniensis* (CD36), and *Candida glabrata* (CBS138), we used the sequenced strains SC5314, CD36, and CBS138/ATCC 2001, respectively. SC5314 was isolated from a patient with disseminated candidiasis prior to 1968 (89–92), CD36 was isolated from the mouth of an Irish HIV patient between 1988 and 1994 (93), and CBS138 is listed as coming from a fecal sample (https://www.atcc.org/products/all/2001.aspx#history). Unless otherwise noted, assays used the same three *C. auris* isolates (AR-384/MLY1540, clade III; AR-386/MLY1542, clade IV; AR- 387/MLY1543, clade I) to ensure representation of the three clades most associated with serious infections as well as wide distribution of sensitivities to fluconazole and caspofungin. A fourth isolate, AR-390/MLY1546 from clade I, was included as the second strain, in addition to AR-384/MLY1540, in the experimental evolution experiment.

A full list of oligonucleotides and plasmids used for the construction of strains can be found in File S4. Strain construction took place in the AR-387 and AR-390 *C. auris* strain backgrounds following previously described methods using the hygromycin B resistance selectable marker (94). gRNA were designed using the gRNA selection tool in Benchling with the following parameters: “Single guide”, “Guide Length” of 20, “PAM” of NGG, with the *C. auris* B8441 (AR-387) reference genome. The gRNA fragments were amplified from pCE41 while the Cas9 construct was prepared by digesting pCE38 with the restriction enzyme MssI. The repair template was created by amplifying genomic DNA and amplicons were subsequently stitched together with an additional round of PCR cycling to create full-length repair template integrating the desired mutation or deletion. When constructing a gene deletion strain, a unique 23bp ADDTAG (CGAGACGAGTGCTCGACATGAGG), which includes a 20 bp gRNA recognition sequence and PAM, was inserted at the location of the gene for any subsequent downstream edits at this locus (95, 96). Synonymous and non-synonymous mutations were introduced using the PCR stitching method and the mutations are denoted by lowercase letters in the oligo sequences described in File S4. The PAM (NGG) and gRNA recognition sequence was mutated to ablate further cutting by Cas9. Transformations used a lithium acetate competence/heat shock-based protocol with a 4-hour recovery before plating on YPD+HYG600. Potentially successful transformations were verified by colony PCR. In order to confirm the presence of desired mutations, DNA for sequencing was extracted using a Quick-DNA Fungal/Bacterial Miniprep Kit (Zymo Research D6005) coupled with a Mini-BeadBeater 16 (Biospec Products); bead beating consisted of two 4-minute cycles separated by a five-minute incubation on ice.

### Antifungal Susceptibility Testing

Antifungal susceptibility testing assays were performed as follows. Overnight cultures (3 mL, in test tubes) were started in RPMI-1640 media on a roller drum at 30°C from two- to three-day old colonies grown on YPD agar plates. The following morning, the OD600 of the overnight cultures was determined, cultures were diluted back to OD600 = 0.25 in fresh RPMI-1640, and the diluted cultures were allowed to recover at 30°C for at least three hours. After the recovery growth step, the OD600 of the recovery cultures was determined and the cells were diluted to an OD600 = 0.00357 in fresh RPMI-1640. Adding of 21 µL of the OD600 = 0.00357 resuspension to 54 µL of media/drug mixture in each well resulted in a starting density of OD600 = 0.001 or approximately 1 x 10^4^ cells/mL.

54 µL of media was dispensed into wells (in two 27 µL steps) using a BioMek FX (Beckman-Coulter). Drugs, DMSO loading controls, and other compounds (e.g. metals) were then dispensed into the media using a Labcyte Echo 525. The 21 µL of cell solution was then added to the 54 µL of media/drug mixture using a BioMek FX. Assays were performed in transparent, sterile, flat-bottomed, non-tissue culture treated 384-well microtiter plates (Thermo 242765 or 242757) that were sealed with Breathe-Easy® sealing membranes (Diversified Biotech, BEM-1) immediately following inoculation. Plates were then incubated at 35°C in a humidified incubator (with 0.1% CO2) for 24 hours. After the 24-hour incubation, the absorbance (OD600) was determined on a prewarmed (35°C) Tecan Spark10M, taking one read per well. In all experiments, the percent inhibition was determined by first subtracting the background OD600 (culture wells filled with media only) from the test well OD600, and then normalizing to the average OD600 of untreated (vehicle only) control wells cultured side-by-side with the test wells. The percent inhibition was then calculated using the following equation: % inhibition = 100*(1- (test well OD - background OD)/(untreated control OD - background OD)).

#### Primary Screening

Initial screening was conducted with the three primary *C. auris* strains. All 1,990 screening compounds were evaluated for *in vitro* efficacy at concentrations of 1 and 10 µM. For each of the strains, a single well was evaluated for each compound at each concentration. The efficacy of each compound was determined by comparison to the average of untreated control wells on the same culture plate as the tested compounds. Compounds with greater than 50% inhibition of *C. auris* growth for at least two of the three tested strains and a B-score (a non-control based method for systematic error correction which accounts for position effects, see (50, 51) for further details) greater than 0.1 were selected as hits for further evaluation (86 compounds total).

#### Secondary Drug Screening and IC50 Determinations

The activity of 86 primary hit compounds was validated against the three primary *C. auris* strains with dose-response growth inhibition experiments. Drugs were dispensed into test wells using the Labcyte Echo 525 liquid handler to generate 8-point concentration ranges from 0.3 to 100 µM. All drugs were resuspended in DMSO and compared to appropriate vehicle-only controls. Dose-response experiments were performed with two biological replicates, each consisting of two technical replicates. Half maximal inhibitory concentrations (IC50s) were calculated from dose-response curves generated in GraphPad Prism 7 using four parameter logistic regression. Subsequent experiments determined the IC50s of the top 5 hit drugs as well as 4 standard-of-care drugs against 13 different *C. auris* strains, as well as one strain each of *C. albicans*, *C. glabrata*, and *C. dubliniensis* with 22-point dose-response ranges (0.05-2600 µM for fluconazole; 0.001-133.3 µM for caspofungin, miltefosine, and triclabendazole; 0.001-20 µM for posaconazole, amphotericin B, chloroxine, broxyquinoline, and clioquinol) performed as described above with three biological replicates, each consisting of two technical replicates. *Human Cell Toxicity Measurements*

The human cell lines Hep-G2 (liver), HEK-293 (kidney), and HFF-1 (fibroblast) were cultured in Dulbecco’s modified Eagle’s medium (Gibco) containing 10% (vol/vol) fetal bovine serum (Gibco), 2 mM l-glutamine, 100 U/mL penicillin/streptomycin (Gibco), and 10 mM HEPES buffer. For toxicity experiments, all human cell lines were seeded into sterile, opaque 384-well culture plates (Corning 3570) one day prior to drug addition. Drugs were added using the Labcyte Echo 525 liquid handler to generate 22-point dose-response concentration ranges.

Cells were cultured in the presence of drug for 72 hours prior to addition of CellTiter-Glo 2.0 reagent (Promega) and collection of luminescence values in relative luminescence units (RLU) using the Promega GloMax plate reader. The percent viability of each treated culture was calculated using the following equation: % inhibition = 100*(test well RLU)/(untreated control RLU). IC50 values for each drug against human cell lines were calculated as described above. Toxicity experiments were performed with three biological replicates, each consisting of two technical replicates.

#### Drug Susceptibility in Evolved and Genetically Engineered C. auris Strains

*C. auris* strains selected for clioquinol resistance or engineered for specific mutations were generated and cultured as described previously and the IC50 of clioquinol and other drugs was determined as described above using 15-point dose-response ranges (0.1-20 µM for clioquinol; 3.3-1333 µM for fluconazole; 0.03-66.7 µM for broxyquinoline and amphotericin B; 0.003-13.3 µM for posaconazole, voriconazole, and micafungin). All strains were tested simultaneously for direct comparison of drug effects and three biological replicates were performed, each consisting of two technical replicates. Fold-changes in IC50 were calculated relative to the parental strains from which each mutant strain was derived.

#### Structure Activity Relationship (SAR) Assays

A group of 32 commercially available drugs or compounds that are structurally related to clioquinol were selected for SAR experiments. All drugs were resuspended in DMSO and tested for inhibition of the three primary *C. auris* strains with 15-point dose-response ranges from 0.01 to 150 µM. Experiments were performed and IC50 values were calculated as described above with three biological replicates, each consisting of two technical replicates.

#### Metal Supplementation Experiments

Stock 10 mM metal solutions for supplementation experiments were prepared in sterile water as follows: Ca^2+^ solution from calcium chloride (Sigma C79-500); Mg^2+^ solution from magnesium sulfate (Sigma 246972); Fe^2+^ solution from ferrous(II) sulfate (Sigma F8048); Fe^3+^ solution from ferric(III) chloride (Sigma F-2877); Cu^2+^ solution from copper(II) sulfate (Sigma C7631); Mn^2+^ solution from manganese sulfate (Sigma M-1144); Zn^2+^ solution from zinc sulfate (Sigma 96500). For metal supplementation experiments, *C. auris* was grown in RPMI media depleted of divalent metal ions using Chelex 100 resin (Biorad) following the manufacturer’s protocol. Drugs and metals were dispensed into culture wells using the Labcyte Echo 525 liquid handler prior to addition of *C. auris* cells. Ethylenediaminetetraacetic acid (EDTA) and *N*,*N*,*N′*,*N′*-tetrakis(2-pyridinylmethyl)-1,2-ethanediamine (TPEN) were used as control chelating agents for comparison to antifungal drugs. The percent inhibition for each drug/metal condition was calculated as described above. For calculation of clioquinol IC50 in the context of different metal concentrations, 8-point dose-responses ranges from 0.01-10 µM were used. Experiments were performed with three biological replicates, each consisting of two technical replicates.

#### Viability Determination by Plating

Viability assays were performed using the AR-384 strain. In brief, overnight cultures (3 mL, in test tubes) were started in RPMI-1640 media on a roller drum at 30°C from two- to three-day old colonies grown on YPD agar plates. The following morning, the OD600 of the overnight cultures was determined, cultures were diluted back to OD600 = 0.7 in 8 mL fresh RPMI-1640, and the diluted cultures were allowed to recover at 30°C for three hours. After three hours, clioquinol (5 µM) and DMSO controls were added to the cultures which were then incubated overnight on a roller drum at 30°C. As a positive control for cell death, an independent culture was pelleted and resuspended in 70% isopropanol for one hour at room temperature (roughly 20-22°C) with vortexing every 15 minutes. Aliquots were taken from the 22-hour clioquinol and DMSO treated cultures as well as 1 hour isopropanol treated cultures, cells were PBS washed, and preliminary 10x stocks of normalized cell density were created based on OD600. The cell density of each sample was then determined by flow cytometry using a BD Accuri C6 Plus; cell counts were based on the number of cells detected in a 10 µL sample. 1x normalized cell density stocks were then created based on these measurements. The exact cell density of these 1x stocks was then determined by flow cytometry of 10 µL of each solution. Next, both high density (1:20 dilution) and low density (1:200 dilution) stocks were made from the 1x stock and plated (50 µL of high density, 60 µL of low density) on YEPD plates at 30°C. Three low density and two high density plates were used for clioquinol and DMSO samples; two low and one high density plate were used for isopropanol samples. After two days, colony numbers were determined using a Protos 3 (Synbiosis) automated colony counter. The input cell density (cells / µL) and detectable colonies (CFU) were both normalized to their respective DMSO treated samples and normalized viability (relative to the DMSO treated samples) was determined by dividing the normalized CFU for each sample by the normalized input cell density.

#### Iron Supplementation Recovery Assay

Iron supplementation recovery assays were performed on the AR-384 strain and flow cytometry for the iron supplementation recovery assays was performed on the previously described BD Accuri C6 Plus. Overnight cultures (3 mL, in test tubes) were started in RPMI- 1640 media on a roller drum at 30°C from two- to three-day old colonies grown on YPD agar plates. The following morning, the OD600 of the overnight cultures was determined, cultures were diluted back to OD600 = 0.5 in fresh RPMI-1640, and the diluted cultures were allowed to recover at 30°C for three hours at which point 4 µM clioquinol was added to the cultures.

Cultures were then incubated a further 21 hours on a roller drum at 30°C. After the 21-hour incubation, two 1 mL aliquots were pulled from each clioquinol treated sample and either iron (2µM each of iron (II) sulfate and iron (III) chloride) or water (equivalent volumes to the iron solutions) was added. Four clioquinol samples were processed in this way. The density of each culture was then determined by flow cytometry and the strains were incubated on a roller drum at 30°C for 25 hours. At each subsequent time point the cultures were vortexed after which samples were removed and diluted with D-PBS. The cell density of each sample was then determined by flow cytometry; cell counts were based on the number of cells detected in a 10 µL sample.

#### Liquid Resistance Assay

Two overnights of AR-384 and one of AR-390 were started from independent single colonies, the following morning the overnight cultures were diluted back to OD600 = 0.05 in fresh media and allowed to recover for three hours. At this point, clioquinol was added to the cultures at 0.75 µM (AR-384) or 0.4 µM (AR-390) and the cultures were incubated for two days on a roller drum at 30°C. Samples were then diluted back to approximately OD600= 0.01 in the presence of fresh media and drug. The three independent cultures were passaged a further 30 times in this manner with passages occurring every three, rather than every two, days after passage 20 (we note that samples were frozen down after passage 20, passage 21 was started from single colonies on plates made from these frozen stocks). Clioquinol concentrations were slowly increased during the course of the experiment in response to increased growth by strains; see File S5 for the clioquinol concentration present for each passage. In parallel to this experiment, eighteen passages were made of two independent control cultures for each strain where equivalent volumes of DMSO were added at each passage.

#### RT-qPCR Methods

Three independent overnight cultures for each strain were started from independent single colonies on roller drum at 30°C, the following morning the overnight cultures were diluted back to OD600 = 0.35 in 5 mL fresh media and allowed to recover for six hours. We note that clioquinol was not present in the overnight or recovery cultures and as such these samples reflect the basal, as opposed to clioquinol-induced, expression levels. Cultures were then spun down, decanted, and the pellets flash frozen with liquid nitrogen before storage at -80°C. Pellets were thawed and RNA was extracted using the MasterPure Yeast RNA Purification Kit (Lucigen MPY03100) followed by DNase treatment with the DNase TURBO DNA-free kit (Invitrogen AM1907). RNA was diluted 1:50 for RT-qPCR which was conducted using the Luna Universal One-Step RT-qPCR kit (New England Biolabs E3005E) on a C1000 touch thermal Cycler / CFX384 Real-Time System (Biorad). Reactions were performed in Hard-Shell PCR Plates (384- well, thin-wall, Biorad HSP3805) sealed with Microseal® B Adhesive Sealers (Biorad MSB- 1001). Two independent oligonucleotide sets each were used for *CDR1*, *CDR2*, and *MDR1*; one of the *CDR1* oligonucleotide sets was located downstream of the E772* mutation. One oligonucleotide set each was used for the control genes *UBC4* and *ACT1*. Two technical replicates were performed for each oligonucleotide set for each biological replicate. The Cq values for the two technical replicates were averaged for each set, after which the averaged Cq values for the *UBC4* and *ACT1* sets for each biological replicate were then averaged. The ΔCt was then determined for each primer set versus the averaged *UBC4* / *ACT1* Cq, after which the ΔΔCt was calculated versus the parental strain and the fold change determined by taking 2^(- ΔΔCt). The average was then calculated for the two probe sets for each gene within each biological replicate, after which the average and standard deviation were calculated for each gene across the three biological replicates.

#### DNA Sequencing

For endpoint samples of the evolved strains, two cultures were inoculated from independent single colonies (single colony samples) and a third culture was started from the dense portion of the streak on the plate (population samples). For samples from intermediate time points for the evolved strains, single cultures were started from the dense portion of the streak on the plate (population samples). A single culture was inoculated from an independent colony for the parental strains. One culture was inoculated from an independent colony (single colony samples) and a second culture was started from the dense portion of the streak on the plate (population samples) for the DMSO control cultures. Cultures for whole genome DNA sequencing were grown in 8 mL media overnight on a roller drum at 30°C. Prior to harvesting the following morning, a sample was pulled to freeze as a glycerol stock for future use. 6.2 mL of each culture was spun down, decanted, flash frozen in liquid nitrogen, and stored at -80°C.

DNA was extracted using the Quick-DNA Fungal/Bacterial Miniprep Kit (Zymo Research D6005) with two 5-minute cycles on a TissueLyser II (Qiagen) separated by a 5-minute incubation on ice; final elution was in 60 µL water and concentrations were determined on a Nanodrop 2000c (Thermo Scientific). The DNA was diluted to 100 µL at a concentration of 10 ng / µL and then sheared using a Biorupter® Pico (Diagenode) with 13 cycles of 30 s on followed by 30 s off and quantified using an Agilent D1000 ScreenTape (Agilent Technologies); DNA fragment size after this step averaged 240 bp.

Library preparation was performed using the NEBNext® Ultra™ DNA Library Prep Kit for Illumina (E7370), using the 200bp recommended bead volumes for step 3, 5 PCR cycles for step 4, and eluting in water for step 5. See File S6 for a list of the i7_index_RC and i5_index_RC oligonucleotides used for each sample. Eluted libraries for sequencing were then quantified via Qubit, pooled, the pooled mixture quantified via Qubit, and then the pooled libraries were quantified using an Agilent High Sensitivity D1000 ScreenTape (Agilent Technologies); DNA fragment size after this step averaged 360 bp.

Sequencing was performed by the Chan Zuckerberg Biohub Genomics Platform on an Illumina NextSeq 550 using a NextSeq 500/550 v2.5 reagent kit (300 cycles, 150bp paired end read, 12bp index length for reads 1 and 2). The number of reads per sample varied between 8,621,624 and 18,340,887 (File S6). Sequences were aligned to the reference genomes using Bowtie2 with default settings, the overall alignment rate varied between 89.38% and 94.57% (File S6), for an approximate sequencing depth ranging from 95x to 200x with an average of 150x. AR-390 based strains were aligned to B8441 chromosome FASTA and GFF features files from the *Candida* Genome Database (version s01-m01-r-17, dating from 8/8/2021, downloaded on 9/10/2021). AR-384 based strains were aligned to B11221 chromosome FASTA and GFF features files from the *Candida* Genome Database (no version information, dating from 12/17/2019, downloaded on 9/10/2021). Aligned reads were then filtered using Samtools to remove reads with a Cigar Value of “*”. Mutations in genes were identified using Minority Report (97) in Python 2 with the parental AR-384 and AR-390 sequencing reads used as the basis for comparison, the analyze “Copy Number Variants” (CNV) feature was enabled, and the codon table changed to reflect the use of “CTG” as Serine rather than Leucine. For single colony samples, the default settings were used; these settings identified mutations that were present in at least 90 percent of the population. For population samples, copy number variation (CNV) analysis was conducted and the following settings were used: “vp” (minimum_variant_proportion) of 0.1, “wp” (maximum_variant_proportion) of 0.04, “vc” (minimum_variant_counts) of 10, and “wc” (maximum_wildtype_variant_counts) of 50. In other words, a mutation or variant must be present in at least 10 percent of experimental sample reads with a minimum requirement for 10 reads and must be present in no more than four percent of parental sequencing reads with a maximum limit of 50 reads. The Minority Report output files for each sample can be found in File S6. As a parallel approach, variant sequences were identified from the alignment sam files which were filtered for map quality using Samools and BCFtools (98) with the output organized from the VCF files using R (99) with extensive use of *tidyverse* (100). The position of each variant was determined to be within a gene (or in an intergenic region) based on coordinates from GFF files, considering ‘gene’ features, which include introns. The results of this analysis are also included in File S6. Additional details explaining the Minority Report and alternative R-based mutation identification outputs are provided in File S7.

### Data Availability

Data from the Illumina sequences analyzed for this project are available at the NCBI Sequence Read Archive (SRA) as BioProject accession number PRJNA932032 (Biosamples SAMN33143520 through SAMN33143533; SRA Experiments SRR23353631 through SRR23353654).

## Acknowledgements

We thank the Centers for Disease Control and Prevention’s Antibiotic Resistance Isolate Bank and UCSF’s Clinical Laboratories at China Basin for strains. We thank Ananda Mendoza for technical support. This work was supported by NIH grants R01AI049187 (to A.D.J.), R35GM124594 (to C.J.N.), an NIH Fellowship F31DE028488 (to C.L.E), and by the Kamangar family in the form of an endowed chair (to C.J.N.). M.T.L. and J.D. were supported by the Chan Zuckerberg Biohub. The content is the sole responsibility of the authors and does not represent the views of the NIH or other funding agencies. Neither the NIH nor other funding agencies had any role in the design of the study, in the collection, analyses, or interpretation of data, in the writing of the manuscript, or in the decision to publish the results.

## Supplemental Materials

**Figure S1:** C**l**ioquinol **is fungistatic to *C. auris*.** (A) Addition of iron restores growth of Clioquinol treated *C. auris* cells. Following a 21-hour incubation in the presence of 4 µM Clioquinol, either iron (2 µM each of iron (II) sulfate and iron (III) chloride, “+ Iron”, squares and solid line) or an equivalent volume of water (“Control”, circles and dashed line) was added to the cells at the 0-hour timepoint and cells were then incubated on a roller drum at 30°C for 25 hours. Cell counts were determined by flow cytometry at the indicated time points; data represent the average of four independent cultures from the same day, error bars represent the standard deviation. (B) Viability of *C. auris* after 22 hours exposure to clioquinol. Aliquots from 22-hour clioquinol (5 µM) and DMSO or 1 hour isopropanol (70%) treated cultures were washed, plated on YEPD plates, and incubated for two days at 30°C. Input cell density as determined by flow cytometry (cells/µL) and the number of colonies were both normalized to the average of their respective DMSO treated samples; normalized viability relative to the DMSO treated samples was determined by dividing the normalized CFU for each sample by the normalized input cell density. Dots represent individual replicates; the mean and standard deviation are indicated for each.

File S1: Ratio of treated *C. auris* viability relative to untreated (vehicle only) control and B-scores for the 1,990 compound primary screen.

**File S2: Mean IC50 for 86 compounds tested in secondary screen.**

**File S3: Mean IC50 for 32 quinoline derivatives tested in structure-activity relationship (SAR) screen.**

**File S4: Compiled information relating to strains, plasmids, oligonucleotides, and compounds used in this study.** The “Strains” tab contains a list of Fungal Strains used in this study. The “Plasmids” tab contains a list of plasmids used in this study. The “Oligonucleotides” tab contains a list of the oligonucleotides used in this study. The “Candidate Compounds” contains a list of the 86 initial candidate and 32 SAR compounds along with sources and product numbers.

File S5: Clioquinol concentrations used for each passage of the experimental evolution study.

**File S6: Compiled information and results relating to experimental evolution strain sequencing.** The “Notes” Tab gives a brief description of the other tabs in the Excel File. The “Sample Guide” Tab provides, among other information, the name, strain background, culture identification, passage number, and experimental treatment for each sample as well as indicating whether each sample was started from a single colony or a population. The “Index Oligonucleotides” tab contains a list of the i7_index_RC and i5_index_RC oligonucleotides used for each sample. The “Alignment Notes” Tab provides the number of Reads and the Overall Alignment Rate (as a percentage) for each sample. The “Full R Call Set” and “R Call Q>200” Tabs contain the information for the mutations called by the alternative R-based (Samtools, BCFtools, tidyverse) mutation identification method. The “R Call Q>200” contains only mutations that were not present in the Parent samples (“PAR_VAR” is FALSE) or DMSO controls (“DMSO_VAR” is FALSE) with a “QUAL” score of at least 200 and a “READS_2” value of at least 50. The 22 tabs with a “SNP_#_S#” format name contain the Minority Report Outputs for the indicated samples (e.g. SNP_3_S3 is the output for Sample 3_S3, an AR-384 DMSO single cell sample corresponding to culture “E” and passage 18). Additional details about this file, including explanations of specific columns in these output files, can be found in File S7.

File S7: Supplemental text explaining the Minority Report and alternative R-based mutation identification outputs.

